# Increasing tolerance of hospital *Enterococcus faecium* to hand-wash alcohols

**DOI:** 10.1101/053728

**Authors:** Sacha J. Pidot, Wei Gao, Andrew H. Buultjens, Ian R. Monk, Romain Guerillot, Glen P. Carter, Jean Y.H. Lee, Margaret M. C. Lam, M. Lindsay Grayson, Susan A. Ballard, Andrew A. Mahony, Elizabeth A. Grabsch, Despina Kotsanas, Tony M. Korman, Geoffrey W. Coombs, J. Owen Robinson, Anders Gonçalves da Silva, Torsten Seemann, Benjamin P. Howden, Paul D. R. Johnson, Timothy P. Stinear

## Abstract

Alcohol-based hand rubs are international pillars of hospital infection control, restricting transmission of pathogens such as *Staphylococcus aureus*. Despite this success, health care infections caused by *Enterococcus faecium* (Efm) - another multidrug resistant pathogen - are increasing. We tested alcohol tolerance of 139 hospital Efm isolates, obtained between 1997 and 2015 and found Efm post-2010 were 10-fold more tolerant to alcohol killing than older isolates. Using a mouse infection control model, we then showed that alcohol tolerant Efm resisted standard 70% isopropanol surface disinfection and led to gastrointestinal colonization significantly more often than alcohol sensitive Efm. We next looked for bacterial genomic signatures of adaptation. Tolerant Efm have independently accumulated mutations modifying genes involved in carbohydrate uptake and metabolism. Mutagenesis confirmed their roles in isopropanol tolerance. These findings suggest bacterial adaptation and complicate infection control recommendations. Additional policies and procedures to prevent Efm spread are required.

## Introduction

Enterococci are members of the gastrointestinal microbiota with low virulence but they have nevertheless emerged as a major cause of healthcare associated infections (*1*). Enterococci now account for approximately 10% of hospital acquired bacteraemia cases globally, and they are the fourth and fifth leading cause of sepsis in North America and Europe respectively (*2*). Hospital acquired enterococcal infections are difficult to treat because of their intrinsic and acquired resistance to many classes of antibiotics (*3*). The difficulties associated with treatment, coupled with the risk of cross transmission to other patients, make enterococcal infections an increasingly important hospital infection control risk (*4*).

Among the medically important enterococci, *Enterococcus faecium* (Efm) in particular has become a leading cause of nosocomial infections (*5*). *E. faecium* population analysis has revealed the emergence of a rapidly evolving lineage referred to as Clade-A1 and includes clonal complex 17 (CC17), comprising strains associated with hospital infections across five continents (*6*, *7*). These hospital strains are resistant to ampicillin, aminoglycosides and quinolones, and their genomes contain a high number of mobile genetic elements and are enriched for genes encoding altered carbohydrate utilization and transporter proteins that distinguish them from community-acquired and non-pathogenic Efm strains (*6*).

A recent Australia-wide survey demonstrated that Efm caused one third of bacteraemic enterococcal infections, and 90% of these were ampicillin-resistant clonal complex 17 strains, of which 50% were also vancomycin-resistant (*8*). Costs associated with the management of vancomycin resistant enterococci (VRE) colonised patients are high because of the need for isolation rooms, specialised cleaning regimes and the impact on staff, bed flow and other resources. Treatment of invasive VRE infections requires higher-cost antibiotics, with patients experiencing side effects and treatment failure due to further acquired resistance (*8*).

Alcohol-based hand rubs (ABHR) and associated hand hygiene programs are a mainstay of infection control strategies in healthcare facilities worldwide and their introduction is aligned with declines in some hospital-acquired infections, in particular those caused by hospital-adapted multidrug methicillin-resistant *Staphylococcus aureus* (MRSA). The compositions of hand hygiene solutions typically contain at least 70% (v/v) isopropyl or ethyl alcohol (*9*-*11*). The application of ABHR for 30 seconds has better disinfection efficacy than traditional approaches with soap and water, with greater than 3·5 log_10_ reduction in bacterial counts considered effective (*12*). The presence of alcohol in ABHR is responsible for rapid bacterial killing at these concentrations although some species are capable of surviving alcohol exposure at lower concentrations (*9*, *13*). The ability to withstand the addition of a certain percentage of alcohol is referred to as alcohol tolerance, and this phenomenon has been described across several genera (*13*, *14*).

To control VRE many healthcare facilities perform active surveillance cultures (ASC) on all patients and then employ contact precautions that involve the use of gowns, gloves and single room isolation for patients colonised (*15*). However, this approach is expensive and cumbersome, particularly when VRE endemicity is high. Due to the relatively low virulence of VRE, other facilities rely on standard precautions, predominantly ABHR usage, and only selectively perform ASC in high-risk areas such as Haematology and ICU (*15*). At Austin Health and Monash Medical Centre, two university teaching hospitals in Melbourne, Australia, patients are screened for VRE rectal colonization on-admission and weekly for all inpatients in defined high-risk clinical areas. VRE-colonized patients are cohorted and contact precautions (including strict adherence to ABHR guidelines) are used routinely (*16*).

In this current study, motivated by our observation that successive waves of new Efm clones that were driving increasing clinical infection, we commenced an investigation into the tolerance of more recent Efm isolates to the short chain alcohol (isopropyl alcohol) used in ABHR.

## Results

### Increasing isopropanol tolerance among hospital Efm isolates over time

ABHR was systematically introduced to Australian health care facilities beginning at Austin Health in December 2002 (*17*-*19*). One consequence of this changed practice has been the substantial increase in the volume of ABHR products used by institutions. For instance, the volume of ABHR increased at Austin Health from 100L/month in 2001 to 1000L/month in 2015. We tested the hypothesis that Efm isolates are adapting to this changed environment and becoming more tolerant to alcohol exposure than earlier isolates. We assessed the isopropanol tolerance of 139 Efm isolates collected from two major Australian hospitals over 19 years. There was considerable variation in isopropanol tolerance, with a range of 4·7-log_10_ between isolates. These differences were independent of Efm sequence type (Data file S1), but we noticed that later isolates were more likely to be tolerant to isopropanol killing than earlier isolates (Fig. 1A), an observation that was supported by significantly different population mean tolerance when comparing pre-2004 with post-2009 isolates (0.97-log_10_ mean difference, p<0·0001, Fig. 1A). There was genetic diversity among the Efm population across this time-period (discussed in more detail later) with two dominant CC17 MLST types (ST17, and ST203) that each incrementally displayed increasing isopropanol tolerance (Fig. 1B,C). Isolates representing the most recently emerged clone (ST796) exhibited uniformly high isopropanol tolerance (n=16, median: 1·14-log10 reduction, Data file S1, Fig. 1D). There was no relationship between acquired vancomycin resistance and isopropanol tolerance. Exposure of a selection of Efm isolates to ethanol showed similar tolerance patterns as isopropanol, with ST796 also significantly more ethanol tolerant compared to representatives of the other dominant Efm sequence types (fig. S1).

**Fig. 1.**
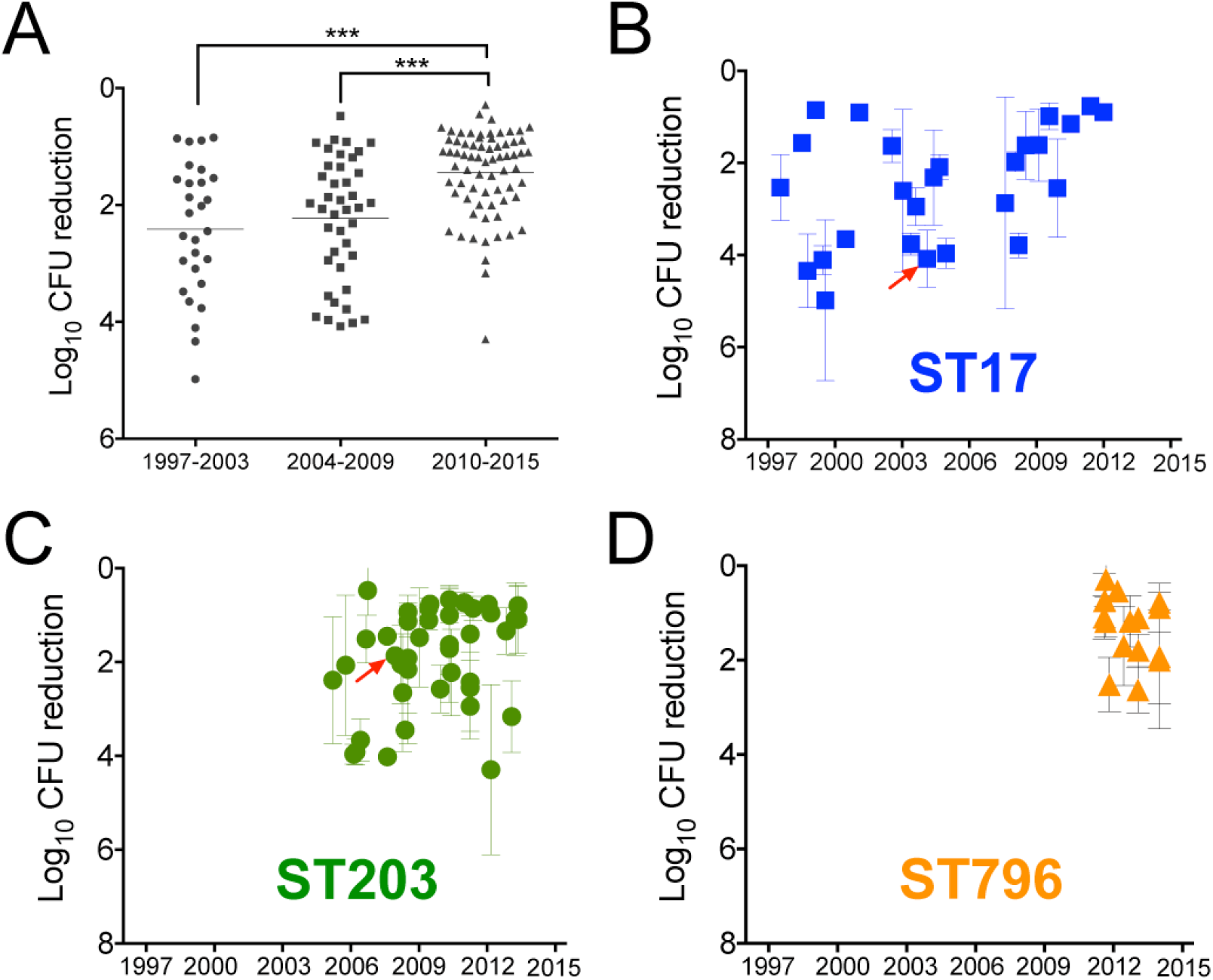
Isopropanol tolerance phenotype variation among 139 Efm isolates over 18 years at two hospitals. (A) Changing isopropanol tolerance between 1998 and 2015. Plotted are the mean log10 CFU reduction values for each Efm isolate obtained after exposure for 5 min to 23% isopropanol (v/v) plotted against specimen collection date and clustered in 5-6 year windows, showing significant tolerance increase of the population-mean over time. Un-paired Mann-Whitney test, two-tailed, p<0.0001. Panels (B), (C), (D) show separately the mean log10 CFU reduction values with range (at least biological triplicates) for each of the three dominant clones. The red arrows indicate isolates used in a previous hand-wash volunteer study (*24*).

### Alcohol tolerance is a clinically relevant phenotype

Our tolerance assay was based on exposure to 23% (v/v) isopropanol, as this concentration provided a discriminating dynamic range among the Efm isolates (refer methods). To assess the clinical relevance of these tolerance differences, we established an Efm contaminated surface transmission model, and compared the impact between two VREfm isolates on transmission of an intervention using 70% isopropanol impregnated surface wipes of a contaminated surface. We employed a mouse gastrointestinal tract (GIT) colonisation model, first establishing that the colonising dose-50 (CD50) among the four Efm isolates was not significantly different (Fig. 2). We selected a 2012, alcohol tolerant isolate (Ef_aus0233, 0.45-log10) and a 1998, reduced tolerance isolate (Ef_aus0004, 4.34-log10). Groups of six BALB/c mice, pre-treated for seven days with vancomycin, were dosed by oral gavage with decreasing doses of each isolate. The CD50 for each isolate was low and not significantly different (Ef_aus0004: 14 CFU, 95% CI 6-36 CFU, compared to Ef_aus0233: 3 CFU, 95% CI 1-6 CFU) (Fig. 2A, B). We then coated the floor of individually vented cages (IVC) with approximately 3 x 106 CFU of each Efm isolate and subjected each cage to a defined disinfection regimen wiping with either water or a 70% v/v isopropanol solution. Groups of six BALB/c mice were then released into the treated IVCs for one hour, before being rehoused in individual cages for seven days and then screened for Efm gastrointestinal colonisation. Across three independent experiments, we then assessed the percentage of mice from each experiment colonised by Efm. The alcohol tolerant Efm isolate (Ef_aus0233) was significantly better able to withstand the 70% isopropanol disinfection and colonise the mouse GIT than the more alcohol susceptible, Ef_aus0004 (p<0.01, Fig. 3C). We then repeated the experiment, but this time using a pair of VSEfm isolates (Ef_aus0026 and Ef_aus0099) also with opposing alcohol tolerance phenotypes, but a much closer genetic association than the first pair (see below for selection criteria). Each isolate had a low CD50 (Ef_aus0026: 19 CFU, 95% CI 9-41 CFU, compared to Ef_aus0099: 12 CFU, 95% CI 3-62 CFU) (Fig. 3B). Ef_aus0099 was 4.4-fold more isopropanol tolerant than Ef_aus0026 with a core genome difference of only 29 SNPs (Data file S2). Across four independent experiments, a significantly greater number mice were colonized by the isopropanol tolerant Efm isolate (Ef_aus0099) than the more susceptible, Ef_aus0026 (p<0.01, Fig. 3C).

**Figure 2:**
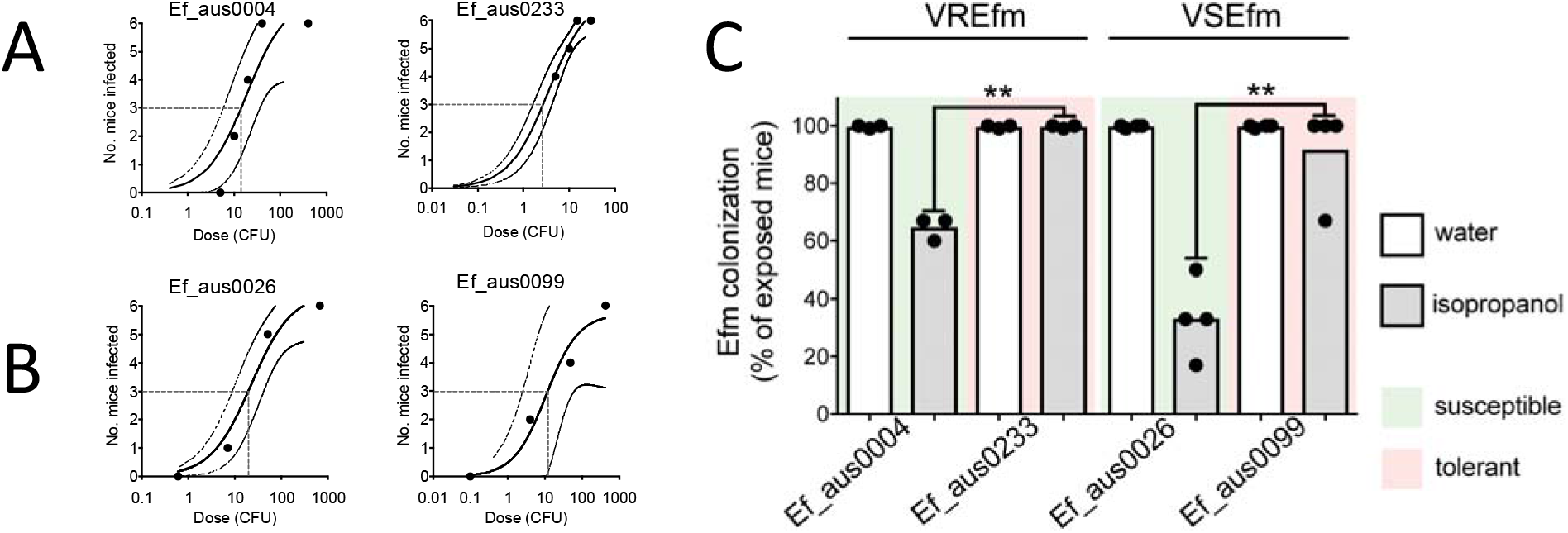
Isopropanol tolerant E. faecium resist disinfection. Transmission model of E. faecium using a murine gastrointestinal colonization model. (A) Establishing the colonizing dose-50 (CD50) for (A) VREfm and (B) VSEfm. Dotted lines indicate CD50. (C) Contaminated cage-floor experiment. Depicted are the percentage of mice colonized with Efm after standardized cage-floor cleaning with either 70% isopropanol or sterile water. Isopropanol tolerant Efm are significantly more likely to be spread. Shown are the results of at least three independent experiments, based on six mice per experiment. The null hypothesis (no difference between sensitive vs tolerant isopropanol) was rejected for p<0.05, unpaired t-test with Welch’s correction.

### Population structure of Efm

To look for signatures of genetic adaptation that associated with alcohol tolerance we first compared the genomes of 129 of the 139 Efm isolates (10 isolates failed sequencing). A high-resolution SNP-based phylogeny was inferred from pairwise core-genome comparisons and Bayesian analysis of population structure (BAPS) that stratified the population into seven distinct genetic groups coinciding with previous MLST designations (Fig. 3A). The population had a substantial pan-genome, comprising 8739 protein coding DNA sequences (CDS) clusters, underscoring the extensive genetic diversity of this Efm population (fig. S2). There was also a temporal pattern to the appearance of each genetic group. Beginning with the previously described displacement of ST17 with ST203 in 2006 through to the emergence of ST796 in 2012, we observed the introduction to the hospital at different times of distinct Efm clones, with each clone exhibiting increasing alcohol tolerance (Fig. 1).

### Identifying bacterial genetic factors linked to alcohol tolerance

High tolerance was observed within distinct Efm lineages, suggesting that multiple genetic events have occurred leading to isopropanol tolerance (Fig. 3). We began the search for the genetic basis of tolerance by evolutionary convergence analysis, to identify regions of the Efm genome that are potentially harbouring genes or mutations linked to alcohol tolerance. We identified pairs of Efm isolates that exhibited greater than 1.5-fold alcohol tolerance difference, and with less than 1,000 core genome SNP differences. With these criteria, there were 19 pairs identified across the 129 isolates (Fig. 3A, Data file S2).

**Fig. 3.**
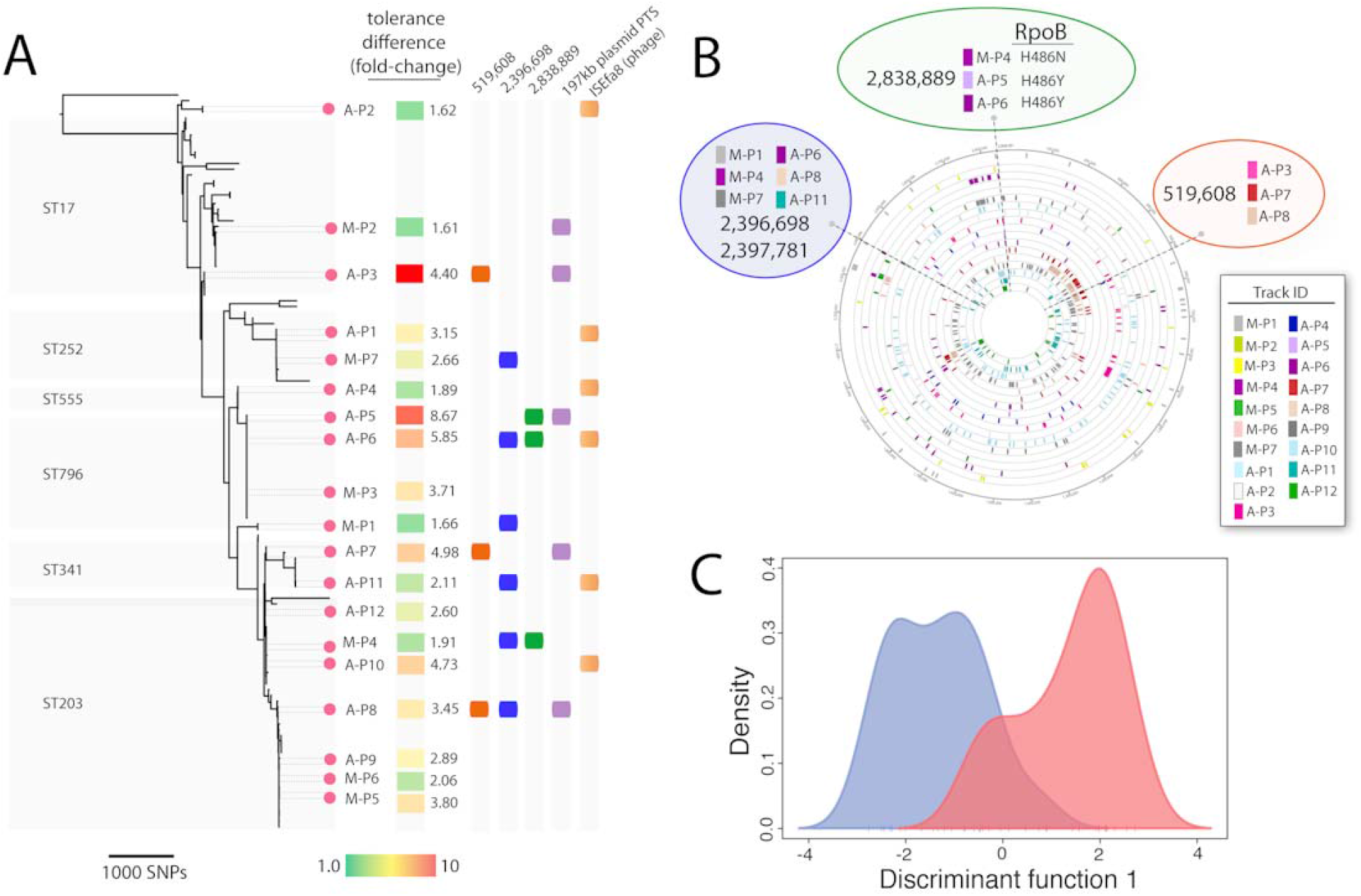
Population structure of E. faecium and identification of tolerant/sensitive pairs. Alcohol tolerance is a polygenic trait. (A) Population structure of the 129 Efm isolates subjected to WGS and alcohol tolerance testing in this study. The phylogeny was inferred using maximum likelihood with RaxML and was based on pairwise alignments of 18,292 core genome SNPs against the Ef_aus0233 reference genome (filtered to remove recombination). Previous MLST designations are indicated. A heat map summary of the log10 kill values is given for each taxon, with green least tolerant and red most tolerant. (B) Analysis of convergent SNP differences among phylogenetically-matched pairs. The prefix ‘A’ indicates Austin Hospital, ‘M’ indicates Monash Medical Centre. Three homoplastic mutations conserved in direction and presence among multiple pairs are highlighted and annotated. (C) Probabilistic separation of sensitive (blue) and tolerant (red) isolates according to a DAPC model built using accessory genome variation.

We then searched for core genome mutations that occurred in different pairs but at the same chromosome nucleotide position and in the same direction of change (i.e. homoplasies). After applying these criteria, three loci were identified (Fig. 3B). One of these loci was the rpoB gene, encoding the beta subunit of RNA polymerase. The H486N/Y RpoB substitution seen in three pairs was associated with reduced alcohol tolerance (Fig. 3). Mutations in this region of rpoB are known to cause resistance to the antibiotic rifampicin, and it is exposure to this drug rather than an evolutionary response to alcohol, that likely selects these mutations. Nevertheless, the rpoB mutations serve as additional support for the approach and its capacity to detect homoplasic mutations associated with a changed alcohol tolerance phenotype. The two additional loci detected spanned an amino acid substitution in a putative symporter in three Efm pairs at chromosome position 519,608 and two mutations in six Efm pairs in a putative phage region (around position 2,396,698) (Fig. 3A,B).

In addition to SNP variations, we also compared the presence/absence patterns of CDS between sensitive and tolerant Efm isolates in each of the 19 pairs. Here, we first used a supervised statistical learning approach called Discriminant Analysis of Principal Components (DAPC) to build a predictive model and identify CDS that contribute to the separation of pairs based on their isopropanol tolerance values. Using only the first 25 principal components, the model showed good separation of sensitive and tolerant isolates, with the resulting loading values used to guide the ranking of CDSs that associated with the alcohol tolerant phenotype (Data file S3). This analysis suggested that there is a genetic basis for the tolerance phenotype, with significant separation of the alcohol tolerant/sensitive populations (Fig. 3C). We then ranked CDS according to (i) their contribution to DAPC separation of the phenotypes, (ii) the frequency of CDS presence/absence among the 19 pairs and (iii) the direction of the CDS presence/absence (i.e. always present in tolerant isolates, Data file S3). This analysis identified two high-scoring loci, a copy of ISEfa8 inserted adjacent to a putative prophage region around chromosome position 953,094 and a 70kb region of a 197kb plasmid that spans CDS encoding several hypothetical proteins, a predicted LPXTG-motif cell wall protein and two PTS systems, named PTS-1 and PTS-2 (Fig. 4A).

### Validation of bacterial genetic factors linked to alcohol tolerance

To test the validity of the predictions based on convergence analysis and DAPC, we used allelic exchange to make targeted mutants in the isopropanol tolerant ST796 reference isolate, Ef_aus0233. Given the reported role of PTS systems in solvent tolerance (*20*), we focused first on one of the plasmid associated PTS regions, deleting a 6.5kb region of PTS-2, a putative glucoside-specific PTS (Fig. 4A). We also made a deletion mutant of the CDS (locus_tag 00501) encoding a putative galactoside symporter (Fig. 4B), where there was a specific V264A aa substitution associated with isopropanol tolerance. An rpoB mutant (H486Y) was also made, as this locus was also identified in the genome convergence analysis and so should present an altered tolerance phenotype. An absence of unintended second-site mutations was confirmed by WGS, and the PTS and 00501 mutations were also repaired. Screening these three mutants and their repaired versions by our isopropanol exposure killing assay showed no change in tolerance (Fig. 4C). To further explore the sensitivity of the two mutants to isopropanol we also conducted growth curve assays in the presence of 3% isopropanol, a concentration we determined provided useful discrimination among our Efm collection. All mutants showed significant increases in their doubling time compared to wild type, a phenotype restored in the repaired mutants (Fig. 4C). The mutants showed no growth defect in the absence of isopropanol (fig. S3). These experiments confirm predictions from convergence testing and DAPC that these loci are involved in promoting isopropanol tolerance. Loss of individual loci however did not impact sensitivity to isopropanol killing, suggesting that isopropanol tolerance is a polygenic phenotype, with multiple genetic changes across different loci likely to have occurred in tolerant Efm strains.

**Fig. 4.**
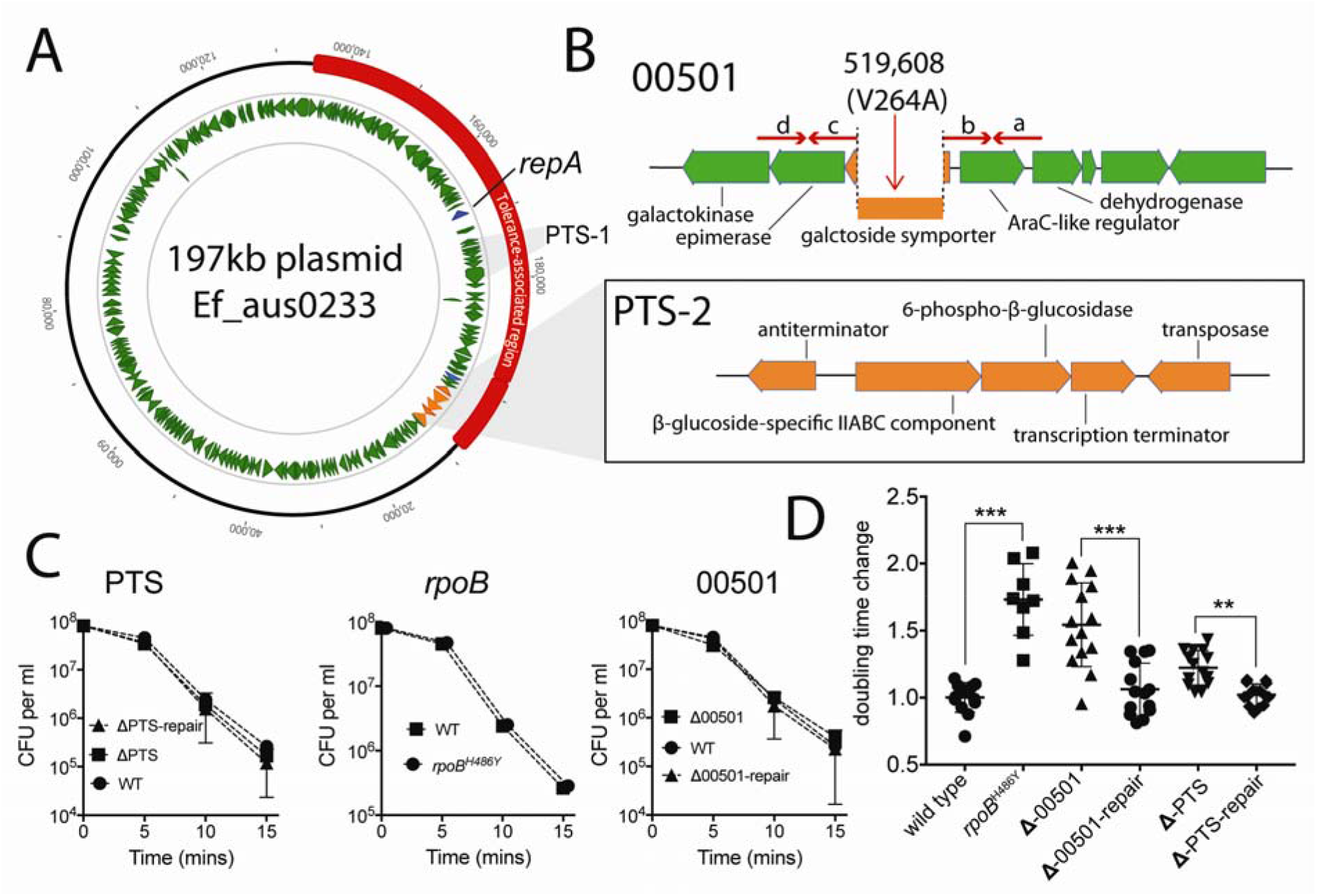
Functional confirmation of genes associated with isopropanol tolerance. (A) Map of the 197kb Efm plasmid, showing the 70kb region associated with isopropanol tolerance (red) and the two PTS loci, including 6.3kb PTS-2 locus (orange) deleted by allelic exchange in the ST796 reference strain Ef_aus0233. (B) Layout of the region around Ef_aus0233 chromosome position 519,608, showing the region deleted by allelic exchange in 00501, encoding a putative symporter. Red arrows indicate position of primers used to generate the recombination substrate for mutant construction (Primer positions: a: 521,394 - 521,371; b: 520,396 -520,420; c: 518,946 - 518,918; d: 517,961 - 517,985). (C) Impact of the mutations on Ef_aus0233 to isopropanol exposure. Shown are the means and SD for biological triplicate experiments, with no difference between mutants and wild type. (D) Growth phenotypes in the presence of 3 % (v/v) isopropanol of the 197kb glucoside ΔPTS locus, rpoBH486Y and Δ00501-symporter mutants. Shown is fold change difference in doubling time for each mutant compared to WT. Depicted also are the phenotypes relative to WT of the repaired mutants. The null hypothesis (no difference between mean doubling time differences of mutant and repaired mutant or WT) was rejected for p<0.01, unpaired Mann-Whitney test. Error bars depict standard deviation. All data points are shown for at least 3x biological duplicates and minimum of 3x technical replicates for each condition.

## Discussion

In 2005, we published a 3-year study describing a progressive decline in rates of hospital acquired methicillin resistant S. aureus and Gram-negative infections following the introduction and promotion of ABHR (*21*). Similar programs were progressively rolled out to all major hospitals in Australia and compliance with ABHR is now a nationally reportable key performance indicator (*19*). The 2015 Australian National Hand Hygiene program report shows increasing and high (>80%) compliance rates in health care facilities across the country (www.hha.org.au) and staphylococcal infection rates have declined nationally (*18*, *22*). However, coincident with the introduction of ABHR and high compliance, there has been a paradoxical nationwide increase in VREfm and vancomycin susceptible *Enterococcus faecium* (VSEfm) infections (*23*).

In this study, we have shown recent clinical Efm isolates were significantly more alcohol-tolerant than their predecessors, and using our *in vitro* alcohol tolerance assay, date of isolation rather than genotype is a better predictor of Efm survival. To obtain a practical dynamic range and allow meaningful comparison between isolates, the tolerance assay used concentrations of alcohol lower than the usual 70% v/v of most ABHR products. However, with our mouse gut colonization model we were able to demonstrate that differences detected by this *in vitro* assay translated to an increased likelihood of transmission for tolerant strains when subject to a full 70% isopropanol surface disinfection intervention (Fig. 2). As tolerance increases, we hypothesise that there will be skin surfaces in contact with ABHR or inanimate surfaces in contact with other alcohol-based cleaning agents that do not receive the maximum biocide concentration or contact time required for effective bacterial killing. This idea is supported by our own previous clinical research using full concentration ABHR in 20 human volunteers and two strains of VREfm (one ST17, one ST203, Fig. 1B, C), and where we identified a mean 3·6-log_10_ reduction in VREfm on the hands of test subjects, but very large inter-subject variance (*24*). For two volunteers, the reduction of VREfm was less than 1·6-log_10_, suggesting that some host factors might not only result in VREfm containment failure (or even “super-spreading”), but also enhance the clinical likelihood for selection of Efm alcohol tolerance (*24*).

Until now there has been no assessment of alcohol tolerance in clinical *E. faecium*, but there has been growing interest in tolerance to other biocides such as chlorhexidine, a second active agent sometimes added to ABHR products (*25*, *26*), including attempts to identify tolerance mechanisms through mutagenesis screens that have pinpointed a specific two-component regulator (*27*). Alcohol tolerance has been reported in other clinically relevant bacteria. For example, studies have reported the enhanced growth of *Acinetobacter baumannii* when exposed to low, non-lethal concentrations of alcohol and ABHR, and increased pathogenicity following the addition of ethanol (*14*, *28*).

Research on alcohol tolerance mechanisms employed by enterococci is scant. Data in this field has been largely derived from studies of Gram-positive bacteria associated with spoilage of sake, in particular the lactic acid bacteria that are known to survive and grow in ethanol concentrations of greater than 18% (v/v) (*29*). The increase in tolerance over time displayed by isolates in our study is consistent with the accumulation of mutations and genes that have shifted the phenotype. Stepwise alcohol adaptation has been observed in laboratory experiments with a related Gram-positive bacterium, *Clostridium thermocellum*, that eventually tolerated up to 8% (w/v) ethanol (*30*). For bacteria in general, short chain alcohols such as ethanol and isopropanol are thought to kill by disrupting membrane functions (*31*, *32*). The penetration of ethanol into the hydrocarbon components of bacterial phospholipid bilayers causes the rapid release of intracellular components and disorganisation of membranes (*33*). Metabolic engineering of solvent tolerant bacteria has uncovered major mechanisms of tolerance, showing that membrane transporters are critically important (*31*). For solvents like ethanol and isopropanol, potassium and proton electrochemical membrane gradients are a general mechanism that enhances alcohol resistance tolerance (*34*).

Our phylogenetic convergence and DAPC analyses in genetically independent Efm populations identified changes in several genetic loci likely contributing to alcohol tolerance. Specific mutagenesis for three regions confirmed these predictions, showing that multiple mutations are required and loci involved in carbohydrate transport and metabolism are likely under selection. No one mutation showed a change in a killing assay after exposure to 25% (v/v) isopropanol (Fig. 4C), but significant differences were observed on growth rate in the presence of 3% (v/v) isopropanol. The CDS (00501) encodes a putative major facilitator superfamily (MFS) galactoside symporter and the SNP at position 519,608 (V264A) occurs within one of its 12 transmembrane regions (Fig. 4B). We speculate that mutations such as V264A might help alter the membrane proton gradient to favour a tolerant state (*34*). In Gram-negative bacteria, transport systems are known to be upregulated or required in response to exposure to short-chain alcohols (*35*, *36*) and bacterial MFS transporters specifically such as 00501 are frequently identified in screens for proteins linked to increased solvent tolerance. However, the specific mechanisms by which they promote tolerance are not understood (*31*). The enrichment in alcohol tolerant Efm strains for PTS loci is also noteworthy (Fig. 4). PTS are bacterial phosphoenolpyruvate (PEP) carbohydrate <u>P</u>hospho<u>T</u>ransferase <u>S</u>ystems (*37*). They catalyse the phosphorylation and transport of different carbohydrates into the bacterial cell. However, there is a growing understanding that their various regulatory roles are equally important as their sugar uptake functions (*37*). Interestingly, they have also been implicated in solvent and stress tolerance. In *E. coli*, up regulation of a mannose-specific PTS led to increased organic tolerance to n-hexane exposure (*20*) and in *E. faecalis* PTS loci appear to be important for survival against low pH and oxidative stresses (*38*). It’s also noteworthy that PTS loci are enriched in health care-associated *E. faecium* lineages, with specific systems associated with GIT colonization, biofilm formation, and serum survival (*39*-*42*).

It is possible that the significant positive relationship between time and increasing alcohol tolerance we report here (Fig. 1) is a response of the bacteria to increased exposure to alcohols in ABHR and that the more tolerant strains are able to displace their less alcohol tolerant predecessors. However, it is also conceivable that Efm are responding to another factor. For instance, modified or acquired transport systems might be conferring acid tolerance, leading to improved survival during passage through the gastrointestinal tract. Secondary phenotypes like alcohol tolerance are then co-selected, passenger phenomena, that together multiply the environmental hardiness of the pathogen.

Whatever the drivers, the development of alcohol tolerant strains of Efm has the potential to significantly undermine the effectiveness of ABHR-based standard precautions, and may partly explain the increase in VRE infection that is now widely reported in hospitals in Europe, Asia, the Americas and Australia. ABHR remains an important general primary defence against cross-transmission of most microbial and some viral pathogens in health care settings. In hospitals with endemic VRE, it would seem prudent to optimise adherence to ABHR protocols to ensure adequate exposure times and use of sufficient volumes of ABHR product each time a healthcare worker cleans their hands. In addition, consideration may need to be given to the use of various formulations of ABHR (e.g. foams and gels) since they are known to have variable (generally reduced) efficacy compared to solutions (*43*). Furthermore, extending active surveillance cultures outside high risk areas of the hospital and return to strict contact precautions during outbreaks with new emergent strains of VRE may be required to prevent widespread cross-contamination.

## Materials and Methods

### Bacterial isolates

Data file S1 in the supplementary appendix lists the 139 Efm isolates investigated in this study that were randomly selected within each year from predominantly blood culture isolates obtained at the Austin Hospital and Monash Medical Centre between 1998 and 2015. Isolates were stored at -80°C in glycerol. Sixty-six of the isolates were vancomycin resistant (60 *vanB-*type, 6 *vanA*-type). Some of these isolates have been described in a previous study on the epidemiology of Efm at the hospital between 1998 and 2009 (*16*) and included recently emergent epidemic clones ST203 and ST796. Six ST341, one ST414 and four ST555 isolates from an Australian-wide enterococci sepsis screening program conducted by the Australian Group on Antimicrobial Resistance (AGAR) were also included, as they were noted emergent clones in other Australian states but were only rarely isolated at our hospitals (*44*). Isolates were grown in on brain heart infusion (BHI) media at 37°C unless otherwise stated.

### Alcohol tolerance assays and analysis

In preliminary experiments, various concentrations of alcohol and Efm inoculum sizes were assessed. At ‘full strength’ isopropanol (70%), killing was complete and resulted in greater than 8-log_10_ reductions in broth culture and an inability to detect differences between isolates. However, by lowering the alcohol concentration in a stepwise fashion, we were able to identify a dynamic range in which we observed marked differences in the time-kill curves between isolates. Guided by these experiments and published literature (*45*) we then measured Efm survival after exposure to 23.0% (v/v) isopropanol. Overnight cultures were grown in 10 mL of BHI medium (Difco, BD). After overnight growth, each strain was diluted to an optical density at 600 nm (OD_600nm_) value of 0.5 using PBS. To 1 mL of the diluted culture, either 23% isopropanol (v/v) or 23% PBS was added and samples were vigorously vortexed, followed by a 5-min incubation at room temperature. Immediately prior to sampling, each culture was again vortexed for 5 seconds and samples were serially diluted between 10-1000 fold in 7.5% Tween80 in PBS (v/v) to inactivate alcohol killing and to give a countable number of colonies on each plate (*46*). An automatic spiral plater (Don Whitley) was used to plate 50 µl aliquots of an appropriate dilution of each strain in triplicate onto BHI agar plates. Plates were incubated overnight at 37°C and colonies were counted using an aCOLyte-3 colony counter (Synbiosis). The limit of detection with this technique was 6000 CFU/ml. For later isopropanol tolerance experiments with mutants, the above killing assay was varied slightly such that that 1ml of 32.5% v/v (final concentration of 25% v/v) of isopropanol was added to 300 µL of cells equating to an OD600nm of 1.66 (~8x10^7^ CFU/ml). These experiments were conducted as described above except spot plates (10µL of dilutions in triplicate) were conducted instead of spiral plating plus additional sampling points were added (10 min and 15 min). Biological replicates were performed for each isolate and average CFU values for cultures exposed to isopropanol and those exposed to PBS (as a control) were obtained. From these data, a mean log_10_ CFU reduction was calculated for each isolate by subtracting the log_10_ CFU remaining after exposure to isopropanol from the mean log_10_ CFU of cultures treated with PBS. Differences in population means for Efm isopropanol tolerance were explored using a Mann-Whitney test with a two-tailed P-value. The null hypothesis (no difference between sample means) was rejected for p<0.05. Statistical analyses were performed using GraphPad Prism (v7.0b).

Growth assays in the presence of isopropanol were performed as follows. Single colonies of Efm were grown in BHI media overnight at 37°C with shaking. The bacterial cell culture concentration was then standardised to an optical density at 600 nm (OD600) of 3.5. Cells were diluted 10-fold in BHI and 10 µL inoculated into 190 µl of BHI broth with or without 3% (v/v) isopropanol. Cells were dispensed in 96-well plates, incubated at 37°C with agitation and the OD600 measured every 10 min over 24 hours using EnSight™ Multimode Plate Reader. The maximum doubling time was determined by fitting local regression over intervals of 1 hour on growth curve data points and by taking the maximum value of the fitted derivative using the R package cellGrowth (www.bioconductor.org/packages/release/bioc/html/cellGrowth.html). The growth rate for each bacterial strain was determined from a minimum of three technical replicates for at least three biological triplicate experiments.

### Whole genome sequencing and bioinformatics analyses

Twenty-two of the isolates examined in the current study have been sequenced previously (*47*-*49*). Genomic analysis and comparisons were performed using established bioinformatics methods that involved assessing Efm population structure and defining core and accessory genomes. Whole genome DNA sequences were obtained using either the Illumina HiSeq or MiSeq platforms, with library preparation using Nextera XT (Illumina Inc). Resulting DNA sequence reads and existing sequence reads were analysed as previously described to define a core genome by aligning reads to the 2,888,087 bp ST796 reference chromosome (Genbank: NZ_379 LT598663.1) (*50*) using Snippy v3.1 (https://github.com/tseemann/snippy). The resulting nucleotide multiple alignment file was used as input for Bayesian analysis of population structure using hierBAPS v6.0 (*51*) and phylogenetic inference using RaxML v8.2.11 (*52*). Whole genome alignments generated by Snippy were used for subsequent assessment of recombination using ClonalFrameML (*53*). Pairwise SNP differences were calculated using a custom R script (https://github.com/MDUPHL/pairwise_snp_differences). Genomes for each isolate were also assembled *de novo* using Velvet v1.20.10 (*54*), with the resulting contigs annotated with Prokka v1.10 (*55*). A pan-genome was generated by clustering the translated coding sequences predicted by Prokka using Proteinortho (*56*) and visualized with Fripan (http://drpowell.github.io/FriPan/).

In order to identify potentially causative variants while reducing the impact of lineage specific effects, pairs of Efm isolates that exhibited greater than 1.5-fold alcohol tolerance difference and less than 1,000 core genome SNP differences were examined. With these criteria, 19 pairs were identified across the 129 isolates. Separate core genome comparisons were undertaken for each the pair using Snippy. The resulting gff files of each within-pair comparison were intersected using bedtools v2.26.0 (*57*) and inspected on the Ef_aus0233 chromosome in Geneious Pro (version 8.1.8, Biomatters Ltd. [www.geneious.com]).

The potential role of gene content variation in the alcohol tolerant phenotype was examined by using a supervised probabilistic approach to assess the contributions of gene presence/absence at separating between sensitive and tolerant isolates. Here, an alignment of accessory genome orthologs was used as input for the generation of a Discriminant Analysis of Principal Component (DAPC) model using the R package adegenet v2.0.1 (*58*). DAPC is a linear discriminant analysis (LDA) that accommodates discrete genetic-based predictors by transforming the genetic data into continuous Principal Components (PC) and building predictive classification models. The PCs are used to build discriminant functions (DF) under the constraint that they must minimize within group variance, and maximize variance between groups.

### Mouse models of Efm gut colonization

Animal experimentation adhered to the Australian National Health and Medical Research Council *Code for the Care and Use of Animals for Scientific Purposes* and was approved by and performed in accordance with the University of Melbourne AEC (Application: 1413341.3). Female Balb/C mice (6-8 weeks old) were used to develop the VREfm and VSEfm gut colonization models. For VREfm colonization, mice were provided drinking water *ad libitum* containing 250mg/L vancomycin for 7 days before VREfm exposure. For VSEfm colonization, after dosing with vancomycin as above, mice were then provided drinking water with ampicillin containing (250mg/L) for a further 7 days. Before exposure of the animals to VREfm or VSEfm, fecal pellets were collected from each mouse to check their Efm status. Briefly, at least two fecal pellets from each mouse were collected and cultured in 10 mL tryptone soy broths (TSB) in a 37-degree shaker for overnight. The cultured broths were then inoculated onto VRE-chrome agar plates for VREfm screening or enterococcosel agar plates for enterococci screening. After 1 week of antibiotic pre-treatment, the mice were dosed by oral gavage with a 200 µL volume of Efm. The bacteria were prepared by culturing overnight in TSB at 37°C with shaking. Bacteria were harvested by centrifugation and washed 3x with sterile distilled water and diluted in sterile distilled water as required before use.

The Efm IVC cage cross-contamination assays were performed in a blinded manner. Bacterial suspensions prepared as described above, were normalized to an OD_600_ 0.37 (~1×10^8^ CFU/mL) and diluted with sterile distilled water to ~1×10^6^ CFU/mL. Each cage was then completely flooded with 10 mL of diluted Efm suspension. Seven millilitres of the suspension were removed from the inundated cage floor. The contaminated cages were left in the biosafety cabinet for 1.5 hours to dry. The dried cage floors (150 x 300mm) were wiped with 40 x 40mm sterile filter paper soaked in 850 µL of freshly prepared 70% (v/v) isopropanol in a consistent manner, with 8 vertical wipes and 24 horizontal wipes in one direction using the same surface of the filter. Each wiping movement partially overlapped the previous. After the isopropanol cage floor treatment, six naïve mice were released into the cage for one hour. Each animal was then relocated to a fresh cage, singly housed and provided with appropriate antibiotics in the drinking water. Fecal pellets were collected from each mouse after 7 days to check the Efm colonization status as described above. CD_50_ values were calculated by interpolation using the non-linear regression and curve-fitting functions in GraphPad Prism (v7.0b).

### Allelic exchange mutagenesis in Efm

To delete a plasmid encoded region encoding a PTS system (6.5kb) and a symporter from the chromosome (1kb), first deletion constructs were PCR amplified (Phusion polymerase - New England Bioabs) (Table S1) from Ef_aus0233 genomic DNA. The construct included 1kb of DNA up/downstream of the region to be deleted and was joined by SOE-PCR. Gel extracted amplimers were cloned into pIMAY-Z (*59*) by SLiCE (*60*). Electrocompetent cells of Ef_aus0233 were made using the method of Zhang *et al* (*61*). Purified plasmid (1 µg) was electroporated, with cells selected on BHI agar containing chloramphenicol 10 µg/ml at 30ºC for 2-3 days. Allelic exchange was conducted as described (*59*) except cells were single colony purified twice pre (30ºC) and post (37ºC) integration. While Efm exhibit intrinsic beta-galactosidase activity, cells containing pIMAY-Z could be differentiated from pIMAY-Z cured cells after 24h at 37ºC. To complement the symporter deletion mutant, the wild type allele for the symporter (amplified with the A/D primers and cloned into pIMAY-Z) was recombined into the symporter deletion mutant. All mutants and complemented strains were whole genome sequenced to ensure no secondary mutations cofounded the analysis.

### Isolation of spontaneous rpoB mutants in Ef_aus0233

An overnight BHI culture of Ef_aus0233 was concentrated 10-fold and 100 µL was spread plated onto BHI agar containing 200µg/ml of rifampicin. A total of three potential *rpoB* mutants were screened by Etest for stable rifampicin resistance. All were resistant to above 32 µg/mL rifampicin. The strains were subjected to whole genome sequencing and single mutations were identified in the *rpoB* gene with one mapping to aa position 481, representing the H481Y substitution.

## Acknowledgments

We thank Takehiro Tomita for DNA sequencing support.

## Author contributions

TPS, PDRJ, BPH designed the study. SJP, WG, IRM, RG, GPC, JYHL, MMCL, SAB conducted laboratory experiments. SAB, MLG, AAM, EAG, DK, TMK, GWC, JOR contributed isolates and obtained metadata. TPS, AHB, AGdS, TS performed bioinformatic analyses. TPS, SJP, AHB, MMCL, SAB, MLG, AAM, EAG, GWC, AGdS, TS, BPH, PDRJ drafted the manuscript.

## Data and materials availability

DNA sequence reads are available from Genbank under study accession PRJEB11390.

## SUPPLEMENTARY MATERIALS

**Fig. S1**. Tolerance of *E. faecium* to ethanol exposure

**Fig. S2**. Core and pan genome analysis of 129 *E. faecium* genomes

**Fig. S3**. Growth curves of mutants

**Table S1:** Oligonucleotides used in this study

**Data file S1:** Strain list

**Data file S2:** Pairwise comparisons of high-low alcohol tolerant *E. faecium*

**Data file S3:** DAPC analysis based on ortholog comparisons versus alcohol tolerance

